# Sleep-dependent offline performance gain in visual perceptual learning is consistent with a learning-dependent model

**DOI:** 10.1101/2020.08.16.253260

**Authors:** Masako Tamaki, Yuka Sasaki

**Affiliations:** Department of Cognitive, Linguistic, and Psychological Sciences, Brown University, USA; National Institute of Occupational Safety and Health, Kawasaki, Japan

## Abstract

Are the sleep-dependent offline performance gains of visual perceptual learning (VPL) consistent with a use-dependent or learning-dependent model? Here, we found that a use-dependent model is inconsistent with the offline performance gains in VPL. In two training conditions with matched visual usages, one generated VPL (learning condition), while the other did not (interference condition). The use-dependent model predicts that slow-wave activity (SWA) during posttraining NREM sleep in the trained region increases in both conditions, in correlation with offline performance gains. However, compared with those in the interference condition, sigma activity, not SWA, during NREM sleep and theta activity during REM sleep, source-localized to the trained early visual areas, increased in the learning condition. Sigma activity correlated with offline performance gain. These significant differences in spontaneous activity between the conditions suggest that there is a learning-dependent process during posttraining sleep for the offline performance gains in VPL.

## Introduction

Numerous studies have demonstrated that sleep is beneficial for various types of learning and memory (Bang et al., 2014; Born & Wilhelm, 2012; Gais et al., 2002; Huber et al., 2004; Maquet et al., 2000; McDevitt et al., 2014; Mednick et al., 2003; Stickgold, 2005; Tamaki et al., 2020a; Tamaki et al., 2020b; Walker et al., 2002; Yotsumoto et al., 2009b). Visual perceptual learning (VPL), which is defined as long-term enhanced performance on a visual task (Lu et al., 2011; Sagi, 2011; Sasaki et al., 2010; Shibata et al., 2011; Watanabe et al., 2001), is one type of learning that shows higher performance after sleep than before sleep. Such improvement over sleep in skill learning (Fischer et al., 2002; Karni et al., 1994) is called offline performance gain (Albouy et al., 2013; Korman et al., 2007; Lugassy et al., 2018; Tamaki et al., 2020b; Vien et al., 2019). However, the underlying mechanism of how sleep results in offline performance gain in learning has been controversial.

There are two opposing models that have been proposed to account for the offline performance gains obtained through sleep in learning: a learning-dependent model and a use-dependent model. A learning-dependent model postulates that offline performance gains are generated by a process that is specifically involved in learning and memory during posttraining sleep. This process may involve synapses or networks that are involved in the acquisition of learning before sleep (Aton et al., 2009; Durkin & Aton, 2016; Rasch & Born, 2013; Stickgold, 2005). Thus, the process is in response to presleep learning. In contrast, a use-dependent model assumes a general process of sleep, such as a homeostatic process. The process simply depends on brain usage during presleep wakefulness, not specifically for learning. However, the process also leads to offline performance gains. For instance, in the synaptic homeostasis hypothesis (SHY) (Tononi & Cirelli, 2003, 2014), one of the use-dependent models, it is postulated that brain usage in prior wakefulness overly potentiates synapses that are downscaled or downselected during sleep, which contributes to offline performance gains on a task (Huber et al., 2004; Tononi & Cirelli, 2003, 2014, 2019). In this case, the process is in response to brain usage, not to presleep learning.

In addition, the range of implicated spontaneous oscillatory activity during posttraining sleep may be different between the learning-dependent models and the use-dependent models. In the learning-dependent models, various spontaneous oscillatory activities are suggested to play a role. Such implicated spontaneous oscillatory activities include slow-wave activity (SWA, 1-4 Hz) (Mascetti et al., 2013; Tamaki et al., 2013) and sleep spindle activity or sigma activity (13-16 Hz) (Bang et al., 2014; Manoach et al., 2016; Tamaki et al., 2019; Wamsley et al., 2012) during NREM sleep, as well as theta activity (5-7 Hz) during REM sleep (Boyce et al., 2016; Nishida et al., 2009). In contrast, the use-dependent models are based on the previous finding that SWA is sensitive to prior sleep deprivation (Borbely, 2001) or brain stimulation during prior wakefulness, including whisker stimulation (Vyazovskiy et al., 2000) and hand stimulation (Kattler et al., 1994). Such use-dependent SWA has been extended to SHY (Tononi & Cirelli, 2003, 2014, 2019). Sigma activity during NREM sleep and theta activity during REM sleep are not particularly assumed to play a major role in the use-dependent model.

In the present study, we examined whether a learning-dependent or use-dependent model (for the latter, SHY) is plausible for the generation of offline performance gains through sleep in VPL. Here, we took advantage of an interference paradigm in VPL in our study. It has been shown that retrograde and anterograde interferences occur in skill learning if two training blocks are performed sequentially without an interval (Faul et al., 2009; A. R. Seitz et al., 2005; Shibata et al., 2017; Yotsumoto et al., 2009a). As a result, no learning is generated after training due to the interference. Based on this observation, we created two conditions, learning and interference conditions. The same number of trials were performed in the learning and interference conditions; thus, visual usage could be considered equivalent between these conditions. However, it is predicted after the equivalent amount of training, learning occurs only in the learning condition and not in the interference condition (**Table 1**).

**Table 1.** The learning vs. interference conditions.

|  | Usage | Learning |
| --- | --- | --- |
| Learning condition | Training | Present |
| Interference condition | Training | Absent |

Critically, the predictions of spontaneous oscillations during posttraining sleep associated with offline performance gains in these conditions are different between the learning-dependent and use-dependent models. A learning-dependent model would predict that the process leading to offline performance gains is present during posttraining sleep only in the learning condition but not in the interference condition. The process should be accompanied by at least one of the 3 bands of spontaneous oscillatory activity in the trained region of the early visual areas. Thus, the strength of the spontaneous oscillations in any of the 3 bands in the early visual areas in the posttraining sleep should be stronger in the learning condition than in the interference condition. Moreover, the stronger oscillations in the learning condition should be correlated with offline performance gain. In contrast, a use-dependent model would predict that the same process, which is represented by increased SWA in response to visual usage prior to sleep, is present in both conditions. Thus, the increased SWA should be found in the trained region in the early visual areas in both conditions and, in particular, should be correlated with offline performance gain in the learning condition. No sigma or theta activity in the trained region in the learning condition should arise or be correlated with offline performance gain. Thus, by examining the spontaneous oscillations during posttraining sleep in the learning and interference conditions, we could assess which of the learning-dependent and use-dependent models is consistent with the offline performance gains observed in VPL.

In the measurement of the strength of spontaneous oscillations, first, retinotopic mapping using functional magnetic resonance imaging (MRI) was conducted to define the trained and untrained regions in the early visual areas (Engel et al., 1994; Yotsumoto et al., 2008). Then, individual structural MRI information was used to source-localize the spontaneous oscillations (Ahveninen et al., 2007; Lin et al., 2004; Tamaki et al., 2013; Tamaki & Sasaki, 2019) to the trained and untrained regions of the early visual areas.

## Results

The subjects were young healthy adults randomly assigned to one of the two conditions, the learning condition (*n*=12, 8 females, mean 23.8 ± 0.80 years old) and the interference condition (*n*=10, 8 females, mean 24.1 ± 1.03 years old). See ***Participants*** in Materials and Methods for more details.

We used the texture discrimination task (TDT), which is one of the standard tasks in VPL (Karni & Sagi, 1991, 1993; Karni et al., 1994; Schwartz et al., 2002; Yotsumoto et al., 2008). Sequential TDT training blocks with orthogonal background orientations are known to cause anterograde and retrograde interference with each other in learning, resulting in no learning (Yotsumoto et al., 2009a), even when the same trained visual field is consistently used across the sequential training sessions.

Two blocks of TDT training were conducted sequentially in both the learning and interference conditions. In the learning condition, the same background orientations (task A) were repeated in the two blocks. In the interference condition, two sequentially presented blocks displayed orthogonal background orientations (tasks A and B). The numbers of trials and the trained visual field quadrant in the training block were the same in these two conditions (see **Supplementary Figure 1** and ***Texture discrimination task*** in Materials and Methods). The only difference in stimuli between the learning and interference conditions was the background orientation in the second training session; it was the same as the first block in the learning condition, whereas it was orthogonal to the first block in the learning condition.

Both the learning and interference conditions included a pretraining test, two blocks of TDT training, and a posttraining test before the 90-min nap, which was followed by the postsleep test (**Figure 1A**). Each of 3 test sessions measured performance for both tasks A and B. In the test sessions, there were 6 stimulus-to-mask onset asynchronies (SOAs) presented (**Supplementary Figure 1** and see Materials and Methods). We obtained the threshold SOA at which subjects marked 80% correct responses in the orientation task as the performance. The performance changes from the pretraining to the posttraining test ([100 × (pretraining − posttraining)/pretraining]) should show the effect of the two blocks of the training session, while those from the posttraining to postsleep test ([100 × (posttraining − postsleep)/posttraining]) should show the offline performance gains generated by sleep, if any (**Figure 1A**).

**Figure 1.**
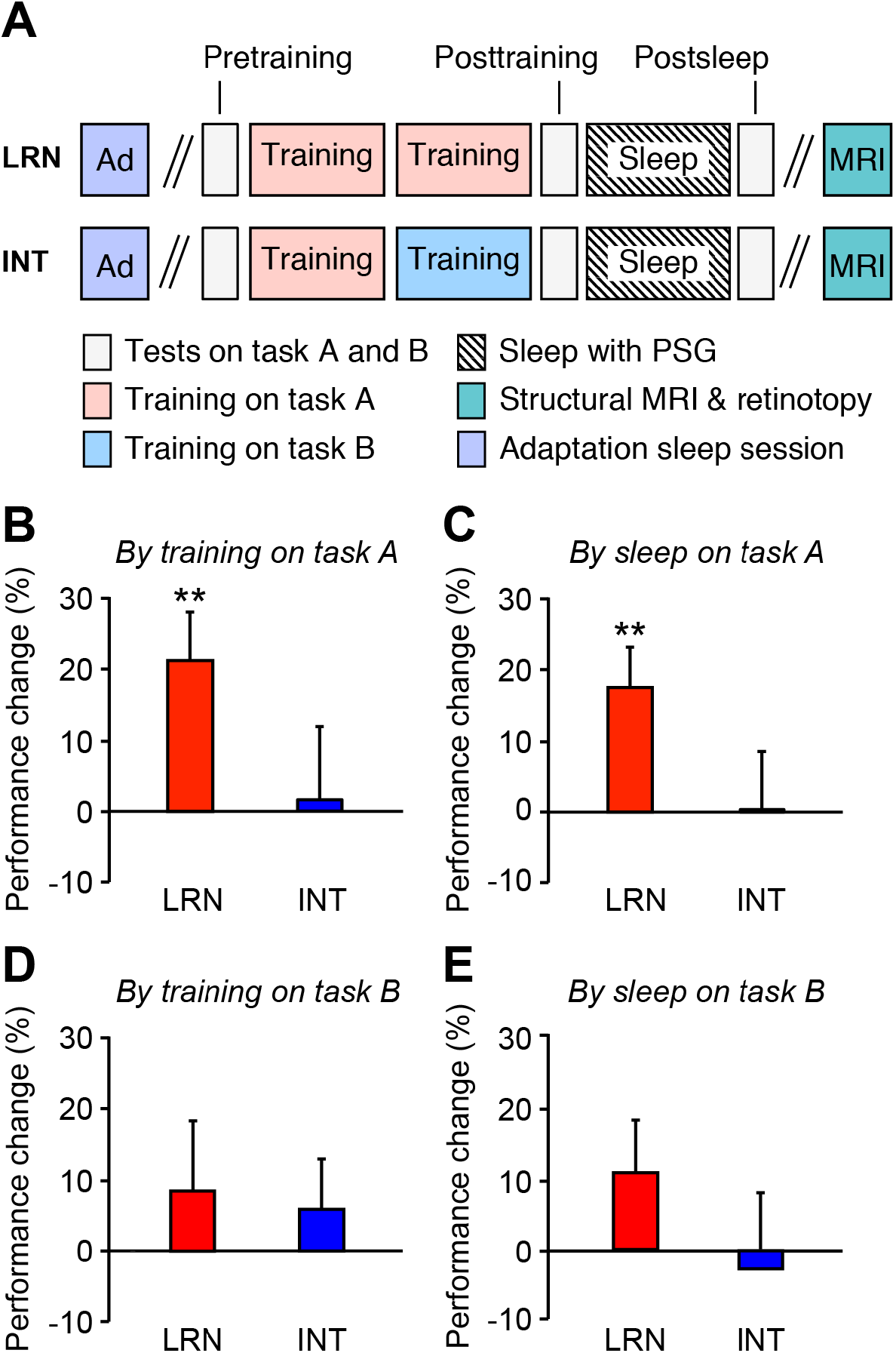
Experimental design and behavioral results. **A**, Experimental design for the learning (LRN) and interference (INT) conditions. In both conditions, an adaptation sleep session (purple) was conducted prior to the main session. An MRI session was conducted in a separate session to obtain structural brain information and retinotopy mapping (green). In the learning condition (LRN), the background line orientation of the TDT stimuli was consistent across the two consecutive training blocks (training on task A, pink). In the interference condition (INT), the background line orientation was switched from the 1st training block (task A, pink) to the 2nd training block (task B, cyan). In both conditions, there were three test sessions: before training (pretraining test), after training and before sleep (posttraining test), and after sleep (postsleep test). In both conditions, a 90-min nap with polysomnography (PSG) occurred for the sleep session. **B**, Mean performance changes (± SEM) from the pretraining to posttraining test sessions for the learning (LRN; red, *n*=12) and interference (INT; blue, *n*=10) conditions on task A. **C**, Mean performance changes (± SEM) from the posttraining to postsleep test sessions for the learning (LRN; red, *n*=12) and interference (INT; blue, *n*=10) conditions on task A. See the main text for details of the statistical results. **D**, Mean performance changes (± SEM) from the pretraining to posttraining test sessions for the learning (LRN; red, *n*=12) and interference (INT; blue, n=10) conditions on task B. **E**, Mean performance changes (± SEM) from the posttraining to postsleep test sessions for the learning (LRN; red, *n*=12) and interference (INT; blue, *n*=10) conditions on task B. See the main text for details of statistical results.

### Performance changes

First, we tested whether the learning condition yielded learning and the interference condition did not. Two-way mixed model ANOVA with a between-subject factor of Condition (learning vs. interference) and a within-subject factor of Test session (posttraining vs. postsleep) was conducted on the performance changes for tasks A and B (**Figure 1B & C**).

For the performance on task A, only the main effect of Condition was significant (*F*(1, 20)=12.24, *p*=0.002). Neither the main effect of Test session (*F*(1, 20)=0.07, *p*=0.790) nor the Condition x Test session interaction (*F*(1, 20)=0.02, *p*=0.902) was significant. For the posttraining test session (**Figure 1B**), a significant performance change was obtained in the learning condition but not in the interference condition (one-sample *t*-test; learning condition, *t*(11)=3.13, *p*=0.010, *d*=0.9; interference condition, *t*(9)=0.17, *p*=0.867, with a Bonferroni adjusted alpha level of 0.025 (0.05/2)). For the postsleep test session (**Figure 1C**), the learning condition showed a significant performance change, or offline performance gains, (*t*(11)=3.16, *p*=0.009, *d*=0.9), but the interference condition did not (*t*(9)=0.05, *p*=0.964, with a Bonferroni adjusted alpha level of 0.025 (0.05/2)).

We also confirmed that the performance on task B did not change in either condition (**Figure 1D & E**). Two-way mixed model ANOVA with the between-subject factor of Condition (learning vs. interference) and the within-subject factor of Test session (posttraining vs. postsleep) was conducted on the performance changes on task B. Neither the main effects (Condition, *F*(1, 20)=1.03, *p*=0.321; Test session, *F*(1, 20)=0.11, *p*=0.747) nor their interaction (*F*(1, 20)=0.32, *p*=0.575) was significant. For the posttraining test session, no significant performance change was obtained in the learning (one sample t-test, *t*(11)=0.86, *p=*0.411) or in the interference conditions (one sample t-test, *t*(9)=0.77, *p*=0.461). For the postsleep test session, no significant performance change was obtained in the learning (one sample t-test, *t*(11)=1.51, *p*=0.160) or in the interference conditions (one sample t-test, *t*(9)=0.28, *p*=0.787).

These results indicate that our experimental design worked as expected: while the same number of trials were performed during training in both the learning and interference conditions, the effects of training and offline performance gain on task A occurred only in the learning condition but not in the interference condition. No learning on task B occurred in either the learning condition or the interference condition.

### Spontaneous brain activity

We measured SWA and sigma activity during NREM sleep and theta activity during REM sleep in the trained and untrained regions in the early visual areas (V1/V2), which were retinotopically identified (Engel et al., 1994; Yotsumoto et al., 2008), as the early visual areas have been indicated to be involved in learning TDT (Karni & Sagi, 1991; Schwartz et al., 2002; Walker et al., 2005; Yotsumoto et al., 2008). Moreover, since the TDT shows a retinotopically specific learning effect (Karni & Sagi, 1991; Schwartz et al., 2002; Yotsumoto et al., 2008), the trained and untrained regions in the early visual areas corresponded contralaterally to the trained and untrained visual field, respectively, were localized. The untrained region served as a control region. Then, we calculated the trained area activity by subtracting the oscillatory activity in the untrained region from the activity in the trained region separately for the SWA and the sigma and theta bands. See ***Source localization of EEG*** in the Materials and Methods for details about the localization of the early visual areas.

First, we tested whether the 3 bands of trained area activity were significantly different between the learning and interference conditions by one-way MANOVA. We used MANOVA to protect against inflating the type I error rate in the post hoc comparisons. After we confirmed that the Box’s M value of 13.054 was not significant (*p*=0.100), suggesting equality of covariance, a statistically significant MANOVA effect was obtained (Pillai’s Trace=0.617, *F*(3,16)=8.608, *p*=0.001, partial η^2^=0.617). Note that only 20 subjects’ data out of 22 were included in this analysis because two subjects did not show REM sleep in the learning condition and were thus excluded.

Next, the results of follow-up one-way ANOVA showed that SWA was not significantly different between the conditions (*F*(1,18)=0.99, *p*=0.333), whereas the sigma (*F*(1,18)=27.23, *p*<0.001, partial η^2=^0.602) and theta (*F*(1,18)=14.48, *p*=0.001, partial η^2^=0.446) activities were significantly different between the conditions.

As noted above, two subjects’ data in the learning condition were removed, as they did not have theta activity data during REM sleep in the MANOVA. Because they had SWA and sigma data during NREM sleep, however, in a supplemental analysis, we included these subjects’ SWA and sigma activity during NREM sleep and tested whether they were significantly different between the two conditions again by one-way ANOVA. The overall statistical results were the same in essence. SWA was not significantly different between the conditions (*F*(1,20)=1.26, *p*=0.274), whereas sigma activity was significantly different between the conditions (*F*(1,20)=22.13, *p*<0.001, partial η^2=^0.525).

Additionally, one sample *t*-tests showed that the sigma (*t*(11)=6.39, *p*<0.001, *d*=1.8, with a Bonferroni adjusted alpha level of 0.017 (0.05/3)) and theta (*t*(9)=3.86, *p*=0.004, *d*=1.2, with a Bonferroni adjusted alpha level of 0.017 (0.05/3)) activities but not SWA (*t*(11)=0.48, *p*=0.639) were significantly different from zero in the learning condition. The results indicate that the spontaneous oscillatory activity in the sigma and theta bands but not SWA was increased in the trained regions of the early visual areas in the learning condition.

Second, we tested whether the sigma and theta trained area activities were correlated with offline performance gain in the learning condition. Both were significantly correlated with offline performance gain, but the latter did not survive the Bonferroni correction for multiple comparisons (sigma activity in **Figure 2C**; *r*=0.72, *p*=0.009, *n*=12, with a Bonferroni adjusted alpha level of 0.017 (0.05/3); no outlier detected by Grubbs’ test; theta activity in **Figure 2D**; *r*=0.70, *p*=0.024, *n*=10, not significant with a Bonferroni adjusted alpha level of 0.017 (0.05/3); no outlier detected by Grubbs’ test). SWA was not significantly correlated with offline performance gain (**Figure 2B**; *r*=−0.07, *p*=0.824, *n*=12). We tested whether increasing the sample sizes would make the correlation stronger. We merged all the data from the learning and interference conditions; regardless, SWA still was only weakly correlated with offline performance gain (*r*=0.172, *p*=0.445, *n*=22).

**Figure 2.**
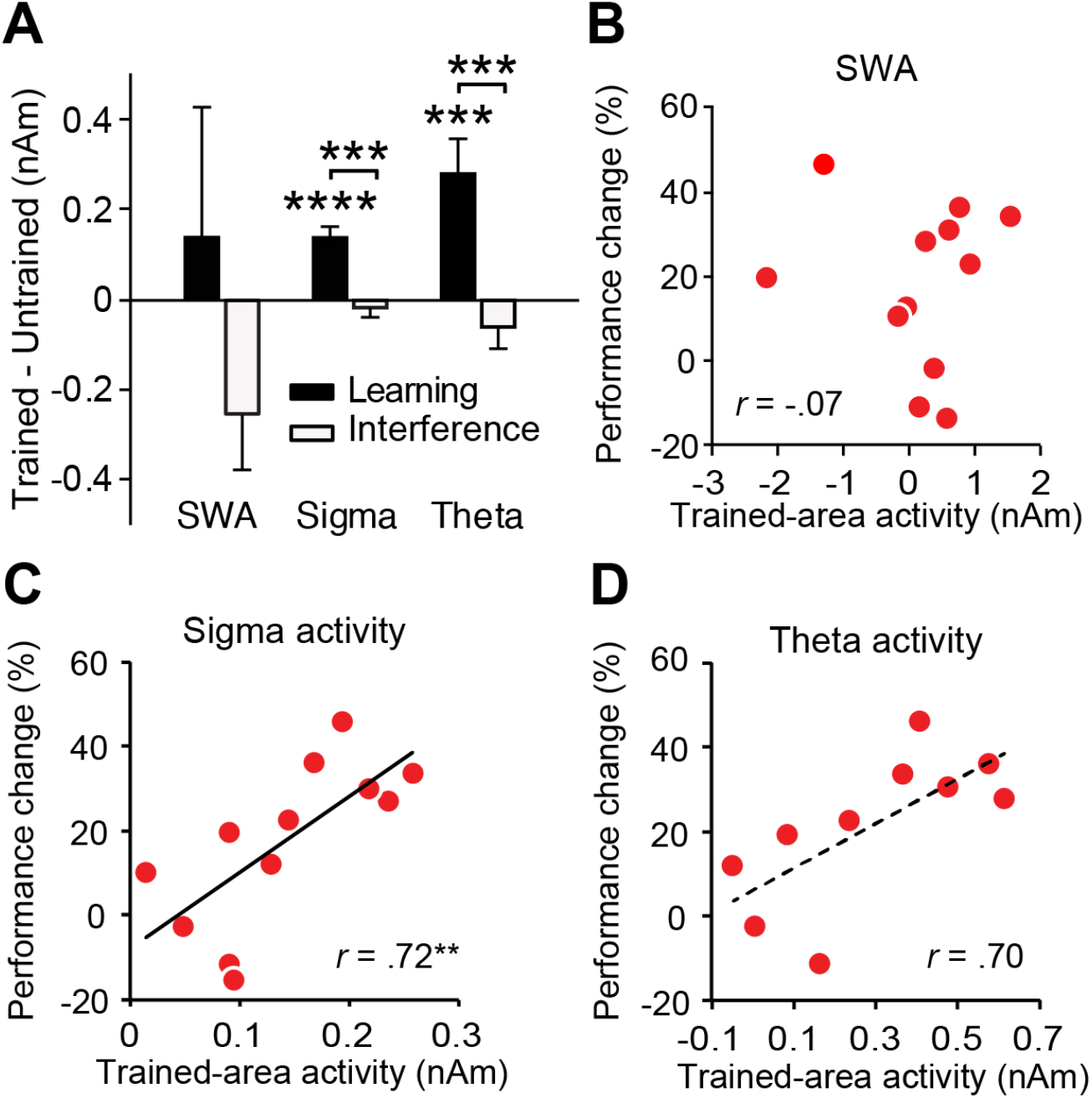
Results for the trained area activities. **A**, Trained area activities during NREM and REM sleep (mean ± SEM) in the early visual areas in the learning (black bars, *n*=12 for SWA and sigma, n=10 for theta) and interference (gray bars, *n*=10) conditions. **B**, Scatter plots for the offline performance gains on task A against trained area SWA during NREM sleep in the learning condition (*n*=12). **C**, Scatter plots for the offline performance gains on task A against sigma trained area activity during NREM sleep in the learning condition (*n*=12). **D**, Scatter plots for the offline performance gains on task A against theta trained area activity during REM sleep in the learning condition (*n*=10). Note that the number of subjects in the learning condition was 12 for **B and C** and 10 for **D**, since 2 out of 12 subjects did not show REM sleep. See the main text for the detailed results of the statistical tests. \*\*\*\**p* < 0.001; \*\*\**p* < 0.005, \*\**p* < 0.01.

These results clearly demonstrate that the strengths of spontaneous oscillations in the early visual areas are consistent with the learning-dependent model: they differed significantly between the learning and interference conditions. In particular, sigma activity during NREM sleep, which was significantly increased retinotopically in the early visual areas in the learning condition, was robustly correlated with offline performance gain.

### Control tests

We conducted several control analyses to rule out that these significant differences in performance and brain activities between the learning and interference conditions were due to factors other than experimental manipulations.

First, we tested whether the initial performance levels were different between the two conditions. We found no significant difference in the pretraining performance between the conditions (task A, *t*(20)=0.56, *p*=0.583; task B, *t*(20)=0.294, *p*=0.772).

Second, we examined whether sleepiness was different between the two conditions. We used two measures to assess sleepiness: the Stanford Sleepiness Scale (SSS) (Hoddes et al., 1973) and a psychomotor vigilance task (PVT) (Dinges & Powell, 1985). The SSS scores subjective sleepiness in scales, while the PVT measures behavioral sleepiness by reaction time. We found no significant differences in sleepiness between the conditions in terms of the SSS scores (Mann-Whitney *U* test, as the sleepiness measures were not normally distributed; pretraining, *U*=55, *p*=0.741; posttraining, *U*=47.5, *p*=0.402; postsleep, *U*=54, *p*=0.674; no corrections for multiple comparisons) or the RTs in the PVT (pretraining, *t*(20)=0.18, *p*=0.856; posttraining, *t*(20)=0.12, *p*=0.906; postsleep, *t*(20)=0.77, *p*=0.452; no corrections for multiple comparisons).

Third, we examined whether any of the sleep parameters was significantly different between the conditions. We did not find any significant differences between the conditions in any of these parameters (see **Table S1**).

Fourth, we tested whether daily sleep habits were different between the conditions. We compared daily sleep quality by the Pittsburg Sleep Quality Index (PSQI) (Buysse et al., 1989) and individual differences in morningness or eveningness by the Morningness-Eveningness Questionnaire (MEQ) (Horne & Ostberg, 1976). There was no significant difference in the PSQI or MEQ scores between the conditions (see **Table S1**).

SOL, sleep-onset latency. SE, sleep efficiency. PSQI, Pittsburg Sleep Quality Index, MEQ, Morningness-Eveningness Questionnaire. Learning condition, *n*=12 (*n*=10 only for REM sleep); Interference condition, *n*=10. Here, we used nonparametric tests because normality was not assumed for most of the parameters (stage W, N1, N3, R, SE%, and MEQ).

The results of the series of control analyses showed that the significant differences in behavioral and spontaneous oscillatory measures between the two conditions were unlikely to be attributed to differences in the initial performance levels, sleepiness, or sleep quality between the conditions.

## Discussion

The results of the present study indicated that the offline performance gains in VPL are in accordance with the learning-dependent model. The strength of spontaneous oscillatory activity during posttraining sleep in the early visual areas was significantly different between the learning and interference conditions, while visual usage was matched between the conditions. The results are consistent with the learning-dependent model, which assumes a learning-specific process during posttraining sleep in response to presleep learning; thus, compared with that in the interference condition, increased spontaneous oscillatory activity was expected in the learning condition.

In contrast, the use-dependent model was not consistent with the present results. The use-dependent model would predict that there would be no significant difference in SWA between the learning and interference conditions. This is what we found; however, an additional two points predicted by the use-dependent model were not found. We did not find a significant increase in SWA in the trained region with respect to that in the untrained region or a significant correlation between SWA in the trained region and the offline performance gains in the learning condition. Because the correlation coefficient was very close to zero, the statistical insignificance in the correlational analysis was due to a very small effect size and unlikely to be due to insufficient statistical power. Thus, use-dependent models, including SHY, are not consistent with offline performance gains, at least in VPL.

In the present study, marked increases were found in different spontaneous brain activities, such as sigma activity during NREM sleep and theta activity during REM sleep, in the early visual areas and were correlated with the offline performance gains in the trained region in the learning condition. However, while the correlation coefficient between the offline performance gains and theta activity during REM sleep was relatively large, it was not as statistically robust as that of sigma activity during NREM sleep. This suggests that sigma activity is primarily involved in offline performance gains, and theta activity may be secondarily involved. A recent study suggests the possibility that theta activity during REM sleep is involved in the stabilization of presleep learning to be resilient to new postsleep learning (Tamaki et al., 2020b). Stabilization of learning is another benefit of sleep (Ellenbogen et al., 2006; Fischer et al., 2002; Gonzalez et al., 2020). In the present study, while we did not measure the degree of stabilization, it is possible that theta activity was involved in the stabilization of offline performance gains.

We measured sigma activity as a substitute to sleep spindles in the present study because sleep spindles around the occipital region have not been clearly defined in the standard manuals (Iber et al., 2007; Rechtschaffen & Kales, 1968). Instead of measuring sleep spindles from the occipital region using arbitrary thresholds for detection, we measured sigma activity, the frequency of which overlaps with that of sleep spindles, source-localized to the early visual areas. This source-localization technique has been shown to be effective in measuring the strength of spontaneous brain activities (Ahveninen et al., 2007). It has been shown that increased sleep spindles are involved in the facilitation of memory and learning (Fogel & Smith, 2006; Tamaki et al., 2008, 2009). Occurrences of sleep spindles may accompany the highest levels of pyramidal cell activity, which may reflect the upregulation of selective synapse formation through local dendritic calcium influx and contribute to learning (Niethard et al., 2017). It is also possible that local sigma activity may induce long-term potentiation (Rosanova & Ulrich, 2005) or replay of circuits or neurons involved in presleep learning (Born & Wilhelm, 2012).

The involvement of brain activity during NREM sleep in the offline performance gains in VPL is consistent with the results of a recent study (Tamaki et al., 2020b). This study demonstrated that the visual plasticity indexed as an excitation and inhibition balance by magnetic resonance spectroscopy was increased during NREM sleep and was correlated with the offline performance gains in VPL (Tamaki et al., 2020b). The results together suggest that after visual training, visual plasticity increases in the early visual areas during posttraining NREM sleep to create a neural environment that promotes sigma activity in the early visual areas, leading to offline performance gains.

One may wonder whether the degree of cortical usage during training was successfully equated between the learning and interference conditions. Such a concern may arise at the orientation column level because the number of trials in both tasks A and B were different between the conditions, although the total number of trials in the training periods including both tasks was equal between the conditions. One orientation was used as the background of the TDT in the learning condition, whereas two orientations were involved in the interference condition. As far as the background orientation is concerned, it is likely that only one orientation column was primarily involved in the learning condition, whereas two were likely activated in the interference condition. However, each orientation task was followed by a mask display that contained multiple orientations in each trial in the TDT. Thus, each trial should excite multiple orientation columns and neurons in the TDT. In addition, spontaneous oscillatory activity measured by EEG is an aggregated sum of the electrical fields generated by many neurons, including multiple orientation columns. It is likely that strengths of spontaneous oscillations originating in different orientation columns may be equally captured by EEG. Thus, we argue that the overall visual usage was equal at the EEG measurement level between the conditions, even though the number of involved orientation columns were different in part of one trial.

Some studies have suggested that SWA is involved in a learning-dependent process for offline performance gains (Diekelmann & Born, 2010; Huber et al., 2004; Tamaki et al., 2013; Tononi & Cirelli, 2003; Wilhelm et al., 2013). However, we did not find that SWA was involved in the learning facilitation process in this study. This is not consistent with the view that SWA is involved in the offline performance gains in VPL. Rather, our finding is consistent with our previous study (Bang et al., 2014) that also showed that the strength of sigma activity in the early visual areas during NREM sleep but not the strength of SWA is involved in the offline performance gains in VPL during sleep. We speculate that the reason why we did not find involvement of SWA in the offline performance gains in VPL is related to the modality of learning and its involved neural circuits. It has been shown that the distribution of SWA is not consistent over brain regions during sleep (Finelli et al., 2001; Iber et al., 2007; Morikawa et al., 1997). SWA appears predominantly over the frontocentral region, with a smaller magnitude over the occipital lobe, which is involved in the current visual task. Thus, one possibility is that SWA is more sensitive to the facilitation of learning or memory, which involves the frontocentral brain regions to a greater degree than the occipital regions. Another possibility, though less likely, is that because the current experiment used naps, the strength of SWA was weaker in general than in nocturnal sleep, making the contribution of SWA to the facilitation process unclear. However, a previous study (Bang et al., 2014) used nocturnal sleep but did not observe contributions of SWA in the facilitation process of VPL. Thus, even in nocturnal sleep, the strength of SWA may not explain the facilitation process of VPL during sleep. Future studies are needed to clarify whether the lack of involvement of SWA in the facilitation process of VPL is due to the modality of learning or insufficient statistical power.

In conclusion, the present study demonstrates that a learning-dependent process can account for the sleep-dependent offline performance gains observed in VPL. At the very least, the offline performance gains in VPL are not consistent with a use-dependent model, such as SHY.

## Materials and Methods

### Participants

To estimate the number of subjects required to reliably indicate the effect of a nap on offline performance gains in VPL, we applied the G*Power program (Faul et al., 2009; Faul et al., 2007)—with the power set to 0.8 and the required significance level α to 0.05 and two-tailed—to our published behavioral data (Tamaki et al., 2019) for a t-test. Furthermore, offline performance gains in VPL were set as the dependent variable and the t-test as the test family in the G*Power program. The result indicated 9 as the minimum sample size. However, we recruited 12 subjects for each condition to ensure that there would be 9 or more data points per condition even when some data had to be dropped due to ineligibility or being an outlier. Thus, a total of 24 young healthy subjects participated in the present study. All subjects gave written informed consent for their participation in the experiments. This study was approved by the institutional review board at Brown University.

Twenty-four subjects were randomly assigned to either the learning (*n*=12) or the interference (*n*=12) condition. Two sets of data were removed from the interference condition. One data set was an outlier (*n*=1, performance change on task A (see ***Texture Discrimination Task***) approximately −125% at the posttraining test, exceeding the 2SD criteria by Grubbs’ test). The other (*n*=1) was due to an ineligibility to participate because of irregular sleep-wake habits. This reduced the total number of subjects in interference condition to 10. The mean ± SEM age was 23.8 ± 0.80 years (8 females) in the learning condition and 24.1 ± 1.03 years (8 females) in the interference condition, with no significant difference between the conditions (independent-samples t-test, *t*(20)=0.21, *p*=0.837).

We conducted a careful screening process for participation eligibility because various factors are known to influence visual sensitivity and sleep stages. The eligibility criteria were as follows. First, subjects were required to have normal or corrected-to-normal vision. Second, all subjects had to have no prior experience in VPL tasks, as prior experience in such tasks may cause a long-term visual sensitivity change (Karni & Sagi, 1991; Lu et al., 2011; Sagi, 2011; Sasaki et al., 2010; Schwartz et al., 2002; A. Seitz & Watanabe, 2005; Yotsumoto et al., 2009b). Third, people who frequently play action video games were excluded because extensive video game playing has been shown to affect visual and attention processing (Berard et al., 2015; Green & Bavelier, 2003; Li et al., 2009). Frequent gamers were defined as those who played action video games at least 5 hours a week for a continuous period of 6 months or more as defined by previous research (Berard et al., 2015; Green & Bavelier, 2003). Fourth, subjects who were included had a regular sleep schedule, i.e., differences in average bedtimes and wake-up times between weekdays and weekends were less than 2 hours, and the average sleep duration regularly ranged from 6 to 9 hours using a sleep-wake habits questionnaire (Tamaki et al., 2020b; Tamaki et al., 2019) and the Munich chronotype questionnaire (Roenneberg et al., 2003). Fifth, based on a self-report questionnaire (Tamaki et al., 2020b; Tamaki et al., 2019), anyone who had a physical or psychiatric disease, was currently under medication, or was suspected to have a sleep disorder was excluded. No subjects were suspected of having insomnia, sleep apnea, restless leg syndrome, sleep walking, narcolepsy, or REM sleep behavior disorder based on a self-reported questionnaire that asked whether the participant had any symptoms known for these sleep disorders (Tamaki et al., 2020a; Tamaki & Sasaki, 2019). Sixth, they had to be aged between 18-30 years.

In addition, as postconsent questionnaires, we conducted the Morningness-Eveningness Questionnaire (MEQ) (Horne & Ostberg, 1976) and the Pittsburg Sleep Quality Index (PSQI) (Buysse et al., 1989) to obtain measures of the subjects’ circadian disposition and their daily sleep quality.

### Experimental design

Starting three days before the onset of the sleep sessions, including the adaptation session (see below) and main sleep session, subjects were instructed to maintain their regular sleep-wake habits, i.e., their daily wake/sleep time and sleep duration, until the study was over. Their sleep-wake habits were monitored by a sleep log for approximately a week prior to the sleep sessions, by which we checked whether they maintained regular sleep-wake habits. On the day before the sleep session, the subjects were instructed to refrain from alcohol consumption, unusually excessive physical exercise, and naps. Caffeine consumption was not allowed on the day of the experiments.

Prior to the main experiment, the subjects participated in an adaptation sleep session. It has been shown that when subjects sleep in a sleep laboratory for the first time, sleep quality is degraded due to the first-night effect (FNE) caused by the presence of a new environment (Agnew et al., 1966; Carskadon & Dement, 1981; Tamaki et al., 2014, 2016; Tamaki et al., 2005; Tamaki & Sasaki, 2019). The adaptation sleep session was thus necessary to mitigate the FNE. During the adaptation sleep session, all electrodes for PSG measurement were attached to the subjects. The subjects napped in the same fashion as they would in the experimental sleep session. The adaptation session was conducted approximately one week before the main experimental sleep session so that any effects due to sleeping during the adaptation nap would not carry over to the experimental sleep session.

On the day of the main experimental sleep session, the subjects arrived at the experimental room at approximately noon (**Figure 1**). Then, electrodes were attached for PSG measurement (see ***PSG measurement***). After the electrodes were attached, behavioral task sessions for the TDT (see ***Texture discrimination task***) were conducted as follows. A short introductory session for the TDT was first conducted to explain how to perform the task and to make sure that all subjects understood the general procedures of the task. After completion of the introductory session, the subjects participated in the pretraining test session of the task to measure each subject’s initial performance level before training. After the pretraining test session, the subjects underwent intensive training on the TDT. The training session consisted of two blocks (tasks A and B), and the stimuli used for these two blocks were different between conditions, although the total number of trials was the same between conditions (see ***Texture discrimination task***). After the training session, the posttraining test session was conducted to measure performance changes from the pretraining to posttraining test. Shortly after completion of the posttraining test session, the room lights were turned off, and the sleep session began at approximately 2 pm. This particular lights-off time was chosen to take advantage of the known “mid-afternoon dip”, which should facilitate the onset of sleep even in subjects who do not customarily nap (Horikawa et al., 2013; Monk et al., 1996; Tamaki et al., 2016). During the sleep session, which lasted 90 min, PSG was measured (see ***PSG measurement***). There was a 30 min break after the sleep session, which was given so that sleep inertia, known to impair performance, could be reduced (Lubin et al., 1976). The time interval between the end of the posttraining test session and the start of the postsleep test session was 120 min (90-min nap + 30-min break). After the 30-min break, the postsleep test session was conducted to measure the changes in performance over the sleep session. There was a short 2-min break between the test and training sessions.

Subjective (SSS) and behavioral sleepiness (PVT) were measured three times prior to each test session (see ***Sleepiness measurement***).

An MRI session (see below) was conducted in a separate session to obtain structural MRI information and for retinotopy mapping (see ***MRI session***).

### Texture discrimination task

The TDT, a standard and widely used VPL task (Karni & Sagi, 1991), was used. The TDT consists of two tasks: an orientation task and a letter task. The orientation task was the main task, whereas the letter task was designed to control the subjects’ fixation (Karni & Sagi, 1991). The orientation task shows retinotopic location specificity where learning occurs only in the trained visual field (Karni & Sagi, 1991; Yotsumoto et al., 2008). In addition, learning is specific to the orientation of background lines (Karni & Sagi, 1991; Yotsumoto et al., 2009a). These findings suggest that early visual areas, in which low-level visual features are processed, are critically involved in improvement in the TDT (Karni & Sagi, 1993; Yotsumoto et al., 2008). Furthermore, the performance on the TDT is known to improve after sleep followed by cortical activation changes in early visual areas during sleep (Bang et al., 2014; Yotsumoto et al., 2009b).

The TDT was conducted in a dimly lit room. The subject’s head and chin were restrained by a chin rest. Visual stimuli were displayed on the computer screen at a viewing distance of 57 cm. Each trial (**Supplementary Figure 1A**) began with the presentation of a fixation point at the center of the screen (1000 ms). Then, a target display was briefly presented (17 ms), followed by a blank screen of varying duration, then by a mask stimulus (100 ms), which was composed of randomly rotated v-shaped patterns. Subjects were instructed to fixate their eyes on the center of the display throughout the stimulus presentation.

The size of the target display was 28°, which contained a 19 × 19 array of background lines. Each background line was jittered by 0.2°. The orientation of the background lines was either horizontal or vertical (**Supplementary Figure 1C**). Each target display had 2 components: a letter (either ‘L’ or ‘T’) presented at the central location of the display (**Supplementary Figure 1D**) and 3 diagonal lines (the target array) presented in a peripheral location within one of the visual field quadrant (the trained visual field quadrant) at a 5–9° eccentricity (**Supplementary Figure 1B**). The target array was aligned either horizontally or vertically on the background in the trained visual field quadrant. The trained visual field quadrant was either the upper left or upper right visual field quadrant, which was randomly assigned to the subjects. The trained visual field quadrant was consistent throughout the trials and sessions for an individual subject. After the mask display, subjects used a keyboard to report whether the central letter was ‘L’ or ‘T’ (letter task) and whether the target array was aligned ‘horizontally’ or ‘vertically’ (orientation task). After the subject’s responses for the letter and orientation tasks were recorded, a feedback sound was delivered to indicate whether the letter task was correct (1000 Hz pure beep) or incorrect (400 Hz pure beep). No feedback was given for responses on the orientation task. The time interval between the target onset and mask is referred to as the stimulus-to-mask onset asynchrony (SOA). The SOA was modulated across trials to control the task difficulty, with a shorter SOA typically resulting in greater task difficulty.

There were 3 test sessions, for which the performance was measured with 120 trials: 6 SOAs (300, 200, 133, 100, 83, 33 ms) each presented 20 times (6 SOAs × 20 trials=120 trials in total). The presentation order of the SOAs was pseudorandomized for each set of 20 trials to reduce the amount of learning and fatigue during the test sessions (Machizawa et al., 2014). Each test session lasted approximately 10 min.

A training session consisted of 2 blocks. The same 6 SOAs were used as in the test sessions, and each SOA was presented in a set of 60 trials (a total of 360 trials) in one block. Thus, the total number of training trials was 720. Each block of the training session started with the longest SOA (300 ms) and then decreased over the trials. Consistent or different orientations of background lines were used for the 2 blocks of a training session, depending on the condition. In the learning condition, the orientations of the background lines were kept consistent between the first and second training blocks (either horizontal or vertical, background A). This design enabled the subjects to show improvements on background A. On the other hand, in the interference condition, the orientations of the background lines were changed orthogonally from the first (horizontal or vertical, background A) to the second (vertical or horizontal, background B) training blocks.

There was a short 10-sec break every 20 trials within the test and training sessions. The training session lasted approximately 60 min, as one block took approximately 30 min.

An introductory session was presented before the pretraining test session. The introductory session was conducted until the subjects reached a certain level of performance. During the introductory session, 3 long SOAs (800 ms, 600 ms, 400 ms) were used. The introductory session started with the longest SOA (800 ms) and was followed by 600-ms and 400-ms SOAs in a blocked fashion, where a set of 20 trials was conducted for each SOA (a total of 60 trials). The introductory session was repeated until the subject could perform the orientation task with at least a 95% accuracy (only one miss out of 20 trials)for the 400-ms SOA trials to ensure that all the subjects knew how to perform the task. The average number (± SEM) of introductory sessions was 1.4 ± 0.26 for the learning condition and 1.9 ± 0.53 for the interference condition (Mann-Whitney U test, *U*=54.5, *p*=0.674). A reminder session was conducted before the postsleep test session to remind the subjects how to perform the task after the 120-min interval. The reminder session had the same 3 sets of SOAs as the introductory session.

The threshold SOAs were obtained for each test session for each subject as follows. The percentage of correct responses for the orientation task was calculated for each SOA. A cumulative Gaussian function was fitted to determine the threshold SOA that corresponded to 80% correct performance using the psignifit toolbox (ver. 2.5.6) in MATLAB (http://bootstrap-software.org/psignifit/ (Wichmann & Hill, 2001)). All trials in which the letter task was incorrect were removed from the calculation of the threshold SOA. The performance changes in the TDT were computed based on the relative changes in the threshold SOAs (ms) between test sessions. The performance change from training (%) was calculated as [100 × (pretraining − posttraining)/(pretraining)]. Similarly, the performance change from posttraining to postsleep (%) was calculated as [100 × (posttraining − postsleep)/(posttraining)].

Stimuli were generated by MATLAB software with psychtoolbox (Brainard, 1997; Pelli, 1997).

### Sleepiness measurement

The Stanford Subjective Sleepiness scale, or SSS rating (Hoddes et al., 1972; Hoddes et al., 1973), ranged from 1 (feeling active, vital, alert, or wide awake) to 7 (no longer fighting sleep, sleep onset soon; having dream-like thoughts). Subjects chose the scale that described their state of sleepiness.

The psychomotor vigilance test, or PVT (Dinges & Powell, 1985), was implemented with the open-source Psychology Experiment Building Language (PEBL) software (Dinges & Powell, 1985). In one trial of the task, after a fixation screen, a target screen was presented in which a red circle appeared in the center. The subjects were required to press the spacebar on a keyboard as quickly as possible upon detection of the circle. The time interval between the fixation and the screen with the red circle varied between 1000–4000 ms. The PVT lasted approximately 2 min. The reaction time (sec) was log-transformed to reduce the skewness of the data (Luce, 1986). The average reaction time was obtained as a measurement of behavioral sleepiness (see ***Control tests*** in Results for sleepiness data).

### PSG measurement

The electrodes for PSG measurement were attached prior to the introductory session, which took approximately 45 min. The PSG measurements consisted of EEG, electrooculogram (EOG), electromyogram (EMG), and electrocardiogram (ECG) measurements. EEG was recorded at 64 scalp sites according to the 10% electrode positions (Sharbrough et al., 1991) using active electrodes (actiCap, Brain Products GmbH, Gilching, Germany) with a standard amplifier (BrainAmp Standard, Brain Products GmbH, Gilching, Germany). The online reference at the Fz electrode was rereferenced to the average of the left (TP9) and right (TP10) mastoids after the recording for analysis. The sampling frequency was 500 Hz, and the impedance was kept below 20 kΩ. The active electrodes included a new type of integrated impedance converter, which allowed transmission of the EEG signal with significantly lower levels of noise than traditional passive electrode systems. The data quality with the active electrodes was as high as 5 kΩ using passive electrodes. The passive electrodes were used for EOG, EMG, and ECG (BrainAmp ExG, Brain Products GmbH, Gilching, Germany). Horizontal EOG was recorded using two electrodes placed at the outer canthi of both eyes. Vertical EOG was measured using 4 electrodes placed 3 cm above and below both eyes. EMG was recorded from the mentum (chin). ECG was recorded from two electrodes placed at the right clavicle and the left rib bone. The impedance was kept at approximately 10 kΩ for the passive electrodes. Brain Vision Recorder software (Brain Products GmbH, Gilching, Germany) was used for recording. The data were filtered between 0.1 and 100 Hz. PSG was recorded in a soundproof and shielded room.

### Sleep-stage scoring and sleep parameters

Sleep stages were scored for every 30-s epoch, following the standard criteria(Iber et al., 2007; Rechtschaffen & Kales, 1968), into stage wakefulness (stage W), nonrapid eye movement (NREM) stage 1 sleep (stage N1), NREM stage 2 sleep (stage N2), NREM stage 3 sleep (stage N3), and REM sleep (stage R).

Standard sleep parameters were obtained (**Table S1**) to indicate the general sleep structure for each experiment. Sleep parameters included the duration of each sleep stage, sleeponset latency (SOL, the latency to the first appearance of stage N2, defined as the sleep onset, from the lights-off), and sleep efficiency (SE, the total percentage spent in sleep)(Iber et al., 2007).

### MRI session

Anatomical MRI data were acquired and used to determine the conductor geometry for the boundary element model (BEM) of the head (Hamalainen & Sarvas, 1989) and to register the EEG sensor locations with the individual subject’s anatomy (Dale et al., 1999; Fischl et al., 1999). Subjects were scanned in a 3T Siemens Trio MR scanner with a 32-channel head coil. Three T1-weighted MR images (MPRAGE; TR=2.531 s, TE=3.28 ms, flip angle=7°, TI=1100 ms, 256 slices, voxel size=1.3 × 1.3 × 1.0 mm^3^) were acquired. Based on these T1-weigted MR images, the cortical surface was inflated for each subject for brain parcellation to localize individual gyri and sulci (Fischl et al., 2004).

Functional MRI data for retinotopic mapping (see below) were acquired to localize the early visual areas using a gradient EPI sequence (TR=2 s, TE=30 ms, Flip Angle=90°). Twenty-five contiguous slices (3 × 3 × 3.5 mm^3^) oriented orthogonal to the calcarine sulcus were acquired covering the occipital to parietotemporal cortices. Data were analyzed with FSFAST and FreeSurfer (http://surfer.nmr.mgh.harvard.edu) software. All functional images were motion corrected (Cox & Jesmanowicz, 1999), spatially smoothed with a Gaussian kernel of 5.0 mm FWHM, and normalized individually across scans. Functional data were registered to the individual reconstructed brains (Dale et al., 1999; Fischl et al., 1999).

A standard retinotopic mapping technique (Engel et al., 1994; Yotsumoto et al., 2008) was employed to determine the trained and untrained regions in the early visual areas for each subject. While the subjects were scanned in the MRI scanner, a flickering checkerboard pattern was presented at the vertical and horizontal meridians with a block design. Based on the blood-oxygen-level-dependent (BOLD) signal contrast between these two meridians, the early visual areas (V1 and V2) were localized for each subject. Additionally, annulus stimuli were used to localize the upper left and upper right visual fields of 5°–9° eccentricity of the early visual areas, which correspond to the size of the trained visual field (Yotsumoto et al., 2009b).

### Source-localization of EEG

To compute the strength of the different brain activities during sleep in the trained and untrained regions in the early visual areas, the EEG data were subjected to Morlet wavelet analysis and source localization using the minimum-norm estimate (MNE) of individual MRI information (see ***MRI session***) as in our previous study (Tamaki & Sasaki, 2019). Morlet wavelet analysis was applied to the raw EEG data (Ahveninen et al., 2007; Lin et al., 2004; Tamaki & Sasaki, 2019) every 30 s to obtain the spectral strength from 1-4 Hz (SWA) and 13-16 Hz (sigma activity) during stages N2 and N3 and from 5-9 Hz (theta activity) during REM sleep. To localize the current sources underlying the EEG signals, the cortically constrained MNE was used on the EEG data using individual anatomical MRI, and the current locations were constrained to the cortical mantle (Ahveninen et al., 2007; Lin et al., 2004; Tamaki & Sasaki, 2019). Information from the EEG sensor locations and the structural MRI segmentation were used to compute the forward solutions for all source locations using a three-layer model of the boundary element method (BEM) (Hamalainen & Sarvas, 1989). The individual forward solutions constituted the rows of the gain (lead-field) matrix. The noise covariance matrix was computed from the raw EEG data for 30 s during wakefulness. These 2 matrices were used to calculate the inverse operator to obtain the estimated source activity during sleep, as a function of time, on a cortical surface (Ahveninen et al., 2007; Lin et al., 2004). The source-localized strength of EEG was averaged across stages N2 and N3 for NREM sleep during the first sleep cycle.

### Statistical analyses

An α level (type I error rate) of 0.05 was set for all statistical analyses. The Shapiro-Wilk test was conducted for all the data to test whether the data were normally distributed. If the data were not normally distributed, nonparametric tests, including the Mann-Whitney *U* test, were used. When normality was not rejected by the Shapiro-Wilk test, Levene’s test was conducted to test for the homogeneity of variance before conducting parametric tests. For parametric tests, MANOVA, ANOVA and the *t*-test were used. Pearson’s correlation was used to test whether there was a significant correlation between the strength of the different brain activities and TDT performance. Bonferroni correction was applied for tests with multiple comparisons. When a statistical test was significant, the effect size was shown.

Statistical tests were conducted with SPSS (ver. 22, IBM Corp.) and MATLAB (R2016a, The MathWorks, Inc.).

## Author Contributions

M.T. and Y.S. designed the research and wrote the manuscript. M.T. performed the experiments and analyzed the data.

## Acknowledgements

This work was supported by NIH R21EY028329.

---

**Figure S1.**
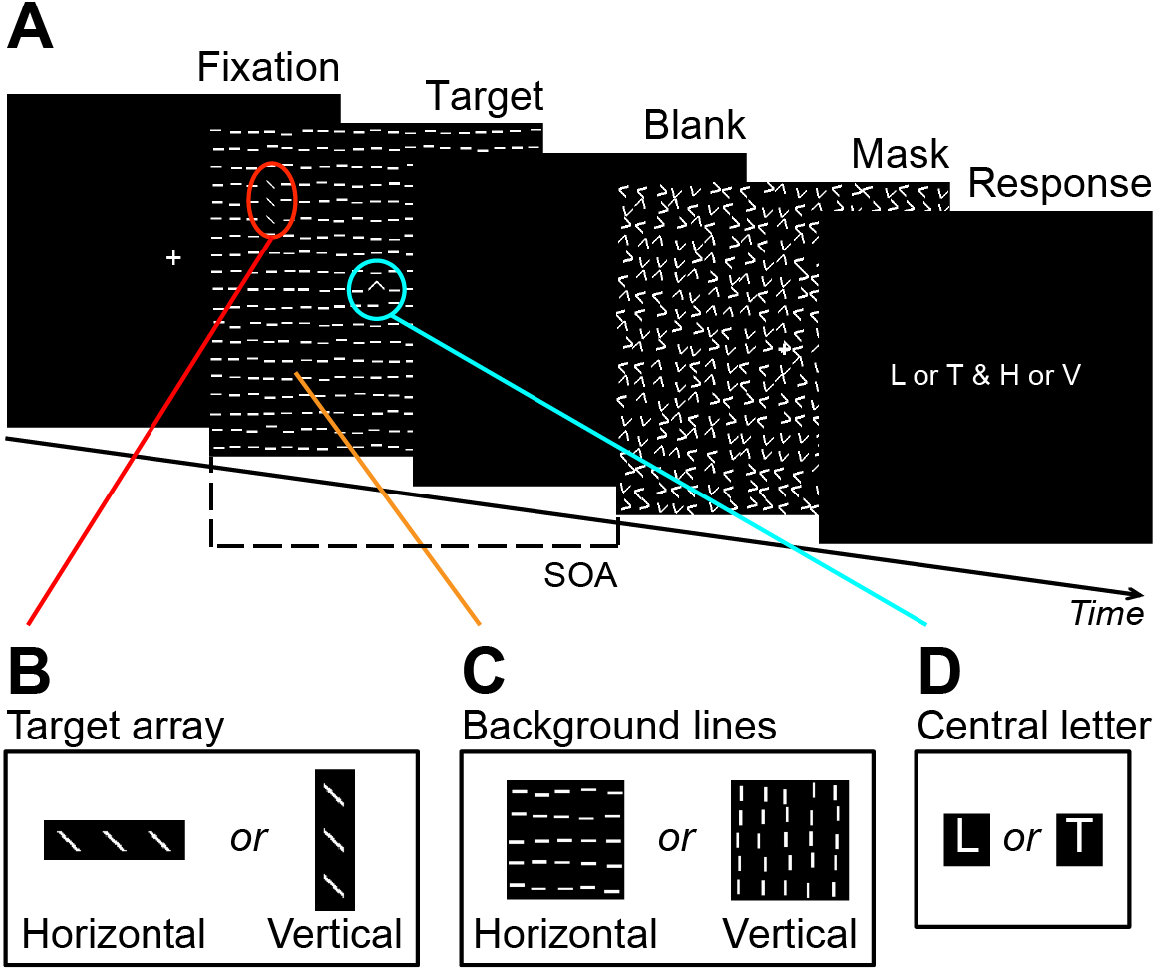
Texture discrimination task (TDT). Stimuli are simplified for visualization purposes. See Materials and Methods for more details. **A.** A sequence of displays in one trial of the TDT. The stimulus-to-mask onset asynchrony (SOA) varied from trial to trial. **B.** Target array. The orientation of an array consisting of three lines was either horizontal or vertical. The array was presented in either the upper left or right visual field quadrant, while the location was consistent throughout task sessions for each subject. **C.** Background lines. The orientation of the background lines was either horizontal or vertical. **D.** Central letter. The purpose of the central letter task was to ensure the subjects’ eye fixation.

**Table S1.** Sleep parameters (mean ± SEM)

|  | Learning condition | Interference condition | Learning vs. Interference |
| --- | --- | --- | --- |
| Stage W (min) | 2.8 ± 1.59 | 1.8 ± 0.63 | $U=58.5$, $p=0.947$ |
| Stage N1 (min) | 8.3 ± 2.28 | 8.8 ± 1.37 | $U=49.0$, $p=0.488$ |
| Stage N2 (min) | 43.0 ± 3.54 | 41.5 ± 3.63 | $U=55.0$, $p=0.767$ |
| Stage N3 (min) | 20.7 ± 4.03 | 13.3 ± 5.21 | $U=38.5$, $p=0.166$ |
| Stage R (min) | 9.3 ± 1.41 | 18.2 ± 4.32 | $U=31.5$, $p=0.173$ |
| SOL (min) | 5.6 ± 0.84 | 7.2 ± 1.23 | $U=50.5$, $p=0.552$ |
| SE (%) | 96.6 ± 1.97 | 97.8 ± 0.83 | $U=59.5$, $p=1.000$ |
| PSQI score | 3.3 ± 0.36 | 3.5 ± 0.34 | $U=56.6$, $p=0.838$ |
| MEQ score | 50.8 ± 2.58 | 51.5 ± 2.46 | $U=56.5$, $p=0.843$ |

SOL, sleep-onset latency. SE, sleep efficiency. PSQI, Pittsburgh Sleep Quality Index, MEQ, Morningness-Eveningness Questionnaire. Learning condition, $n=12$ ( $n=10$ only for REM sleep); Interference condition, $n=10$ . Here, we used nonparametric tests because normality was not assumed for most of the parameters (stage W, N1, N3, R, SE%, and MEQ).

